# Dissimilarity-Overlap Analysis of Replicate Enrichment Communities

**DOI:** 10.1101/2020.02.26.966473

**Authors:** Jean C. C. Vila, Yang-Yu Liu, Alvaro Sanchez

## Abstract

The taxonomic composition of microbial communities can vary substantially between individual hosts and within the same host over time. Efforts to build quantitative and predictive models of microbial population dynamics are underway, but fundamental questions remain. How different are population dynamics in different hosts? Do communities that share the same taxa also exhibit identical dynamics? *In vitro* communities can help establish baseline expectations that are critical towards resolving these questions in natural communities. Here, we applied a recently developed tool, Dissimilarity-Overlap Analysis (DOA), to a set of experimental *in vitro* communities that differed in nutrient composition. The Dissimilarity and Overlap of these communities are negatively correlated in replicate habitats, as one would expect if microbial population dynamics were on average strongly convergent (or “universal”) across these replicate habitats. However, the existence of such a negative correlation does not necessarily imply that population dynamics are always universal in all communities. Even in replicate, identical habitats, two different communities may contain the same set of taxa at different abundances in equilibrium. The formation of alternative states in community assembly is strongly associated with the presence of specific taxa in the communities. Our results benchmark DOA, providing support for some of its core assumptions, and suggest that communities sharing the same taxa and external abiotic factors generally (but not necessarily) have a negative correlation between Dissimilarity and Overlap.

## Introduction

Microorganisms grow and thrive in all habitats throughout the biosphere [1–4]. This includes the human body, where they form rich ecological communities made of large numbers of interacting species [5–8]. The taxonomic composition of these communities can vary substantially between body sites, reflecting their different ecological, physical, and biochemical conditions [9]. Even for the same body site, community composition may vary widely between individuals, as well as within the same individual over time [10, 11]. In order to understand how microbiomes change longitudinally and over the lifespan of an organism, and to design effective strategies that enable us to manipulate microbiomes towards desirable states, it is critical to developing predictive quantitative models of microbial population dynamics [12].

Models of dynamical ecosystems vary in their level of description, which is typically chosen to capture the specific phenomena under study. A detailed population dynamics model of microbial communities would have to include mechanistic microbial interactions (due to cross-feeding [13, 14], direct secretion of substances such as bacteriocins, antibiotics, or extracellular enzymes [15, 16], or competition for the same nutrient), spatial structure of the particular habitat [17], and environment- or host-microbiome interactions. Building such detailed models can be daunting due to (i) a huge number of model parameters which need to be inferred from experimental data; and (ii) many environmental variables (such as the concentrations of bacteriocins and nutrients) are hard to measure in real time.

To avoid those difficulties, an alternative modeling framework focuses on exploring the impact that any given microbial species has on the abundance of other microbial species [18]. In this phenomenological modeling framework, one only needs to consider a simple population dynamics model written as a set of ordinary differential equations (ODEs): d*X*(*t*)/d*t* = *f*(*X*(*t*), Θ). Here, *f* is a nonlinear function characterizing the population dynamics of the microbial community, *X*(*t*) = (*x*_1_(*t*),…, *x_i_*(*t*),…, *x_N_*(*t*)) is an *N*-dimensional vector with *x_i_*(*t*) denoting the abundance of the *i*-th microbial species at time *t*, and Θ captures all the ecological parameters (such as intrinsic growth rates, intra- and inter-species interaction strengths, etc.). Note that those ecological parameters depend on environment- or host-independent factors, such as biochemical processes and microbial metabolic pathways; as well as environment- *or* host-specific ones, such as pH, temperature, nutrient intake, host genetic make-up, etc. Hence, environmental or host factors are not explicitly considered in this modeling framework but are absorbed in the ecological parameters [18].

Generally, the ecological parameters estimated from a given habitat with certain characteristic environmental conditions do not necessarily map to other habitats with different environmental conditions. One can ask, however, whether those parameters (Θ) are strongly similar (“universal”) for microbiomes that assemble in similar habitats. Addressing this fundamental question has important consequences for the applicability and predictive power of quantitative models of microbial community dynamics. If the interaction parameters were highly similar across habitats of a certain type, such as the guts of different human subjects, this will facilitate the development of generic microbiome-based therapeutics. By contrast, if the ecological parameters and microbial dynamics are strongly host-specific, we must design truly personalized interventions, which need to consider not only the highly personalized microbial composition of each individual but also the unique dynamics of the underlying microbial ecosystem.

Directly addressing the above question would require one to infer all of the ecological parameters and fit the population dynamics *f*(*X*(*t*), Θ) from the microbiome data of each local community or host. Doing this for a large collection of communities is both logistically and computationally challenging. Recently, an indirect method called Dissimilarity-Overlap Analysis (DOA) was proposed [18]. DOA relies on two mathematically independent measures between any two local communities: Overlap (*O*), which is defined as half of the sum of relative abundances of the shared species; and Dissimilarity (*D*), which is defined as the divergence between the renormalized abundance profiles of the shared species (Methods) [18]. DOA is based on the following two assumptions. First, the abundance profiles of the microbiome samples represent the steady states *X*\* of the microbial ecosystem and hence the fixed points of the underlying population dynamics that satisfy *f*(*X*\*, Θ)=0. Second, if any two local communities that have the same species collection also have the same abundance profile (steady state), i.e., *O*=1 and *D*=0, then the two communities should share universal microbial dynamics *f*(*X*, Θ) characterized by the same set of ecological parameters Θ. Mathematically, this means that if *X*\* satisfies both *f*(*X*\*, Θ^(1)^)=0 and *f*(*X*\*, Θ^(2)^)=0, given the large number of species and all the other levels of complexity in their interactions (encoded in the highly nonlinear function *f*), we conclude that generically Θ^(1)^=Θ^(2)^. In general, since *D* is mathematically not constrained by any value of *O*>0, any constraints of *D* by *O* observed from real data deserve ecological interpretations. In particular, even if we don’t have any steady state pair satisfying *O*=1 and *D*=0 (which is the typical case for host-associated microbial communities, such as the human gut microbiome, due to highly personalized microbial compositions), often we observe that steady state pairs with higher *O* tend to have lower *D*, i.e., there is a negative slope in the high-overlap region of the Dissimilarity-Overlap Curve (DOC). This particular statistical constraint of *D* by *O* is consistent with the hypothesis of universal dynamics across all samples in the habitats, which is a foundation of DOA [18]. This statistical constraint of *D* by *O* is also consistent with alternative hypotheses, such as communities assembling in environmental gradients, or situations when only a small fraction of the habitats have highly similar interaction parameters [19]. The former is a particularly important scenario, and was recognized in the original study by Bashan et al. In many instances enough is known about the habitats to exclude from the analysis factors that can lead to environmental heterogeneity [18].

A negative slope in the high-overlap region of the DOC has been found in the gut microbiome samples collected from different healthy individuals [18]. Yet, the complete set of selective pressures experienced by microorganisms in the same habitat (e.g., the same body site of different individuals), and their variation across a host group can not be known exactly. Hence, one cannot account for all the potential environmental or host factors that may conceivably influence the microbial communities assembled in the same habitat, and so cannot provide an entirely conclusive answer regarding the universality of the underlying microbial dynamics. In other words, we cannot unambiguously attribute the negative slope of DOC to universal dynamics and completely rule out the alternative explanation of environmental or host factors. A more definitive demonstration would require a comparison between experimental communities assembled in well-controlled replicate habitats to those assembled in non-identical habitats.

Benchmarking DOA against well-controlled *in vitro* communities, ideally assembled in multiple replicates of habitats that are either identical to each other, or different from each other in well-understood ways, would be necessary to understand the limitations and potential of DOA for its application to natural communities [20]. To address this need, here we perform DOA for a large set of *in vitro* communities that meet these requirements: close to three hundred independent enrichment communities assembled in multiple, replicate synthetic habitats on three different limiting nutrient conditions, and assembled to equilibrium under periodic serial dilution cycles [13].

## Results

### Data Set

In a recent study [13], we reported the assembly of a total of 276 enrichment communities in three different synthetic environments: M9 minimal medium with either glucose, citrate or leucine as the only carbon source. These enrichment communities were assembled from twelve different environmental sources (including various soil samples and plant matter collected near Yale University in New Haven, CT). Seven or eight biological replicates of each inoculum were propagated in each of the three nutrient environments, under serial dilution with transfers every 48 hr with a dilution factor of 125x. A diagrammatic summary of the experiment is presented in **Fig. 1A**. As reported in [13], communities were initially very diverse (N=110-1290 unique Exact Sequence Variants, or ESVs). They typically converged to an approximately stable composition (containing N=2-22 ESVs) after 50-60 generations, suggesting that communities were close to a steady state. Metabolic cross-feeding was found to be widespread and critical for the coexistence of multiple species on a single limiting resource [13].

**Fig. 1:**
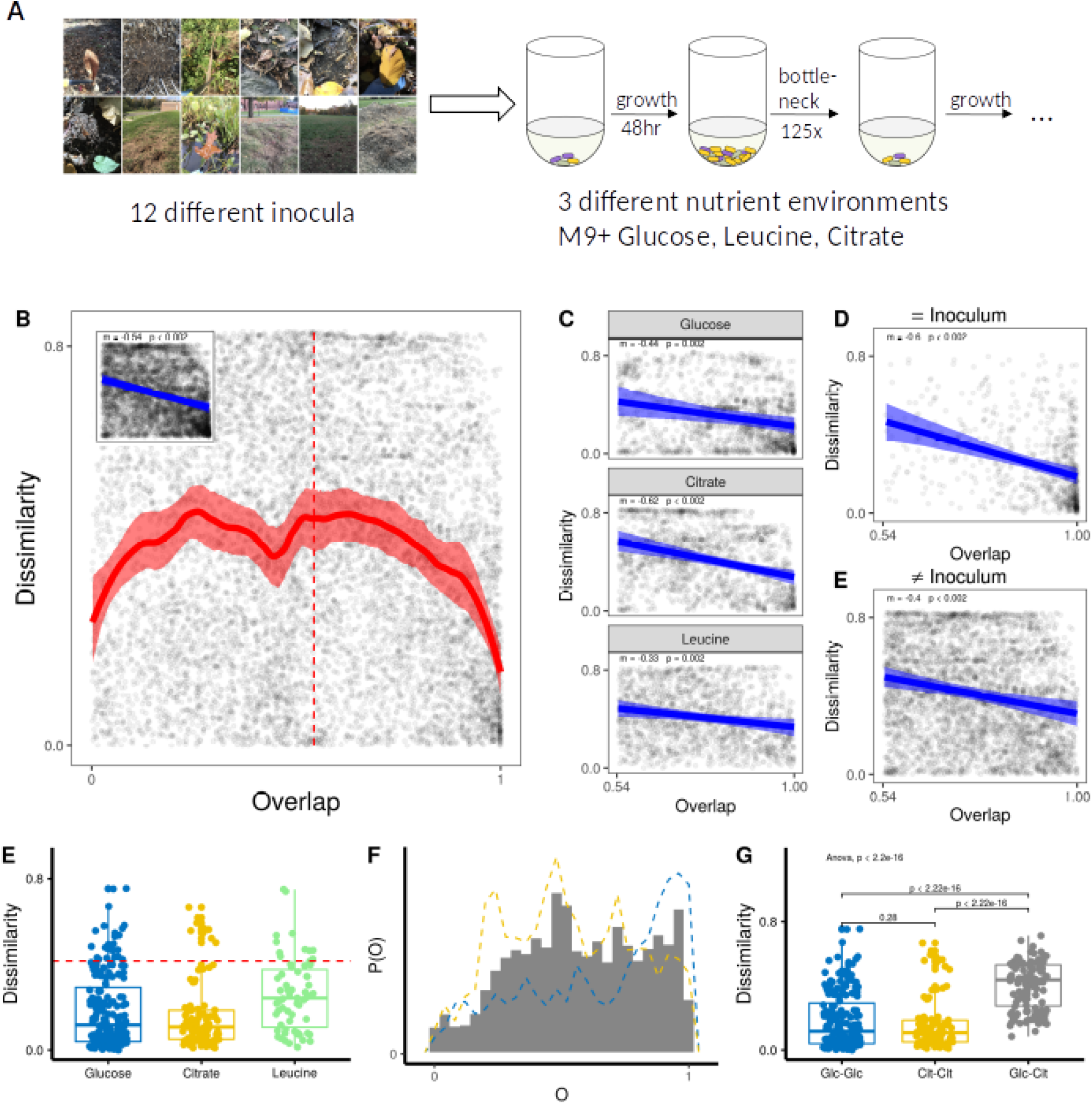
Communities assembled in the same environment show a negative correlation between Dissimilarity and Overlap. (A) Schematic description of the experiments in ref: [13] (B) DOC of all microbial community pairs that have been assembled in the same environment (n = 276 Samples). Shaded regions indicate the 95% confidence interval (methods). The vertical dotted red line represents the median Overlap (0.543). The inset shows a linear regression for communities above the median overlap. We repeat this regression over the same region, subsetting the data to consider (C) each nutrient environments separately; (D) subsets of pairs that have been assembled from the same inoculum (D); subsets of the pairs that have been assembled different inoculum. For each regression, we report m (slope of the linear regression) and a p-value calculated as the fraction of bootstrap realization in which this slope is negative (see methods). (*F*) Distributions for community pairs assembled in the same environment (both in glucose, both in citrate, or both in leucine) with high overlap (O>0.98). The dotted red-line is at half the maximum possible 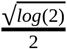. (G) Histogram showing distributions of Overlaps for community pairs where one has been assembled on glucose and the other on citrate. The dotted lines give the frequency polygon for Glucose-Glucose Community pairs and Citrate-Citrate community pairs (Blue and yellow respectively). We use the same binwidth (.04) for both histograms and frequency polygons so the two are comparable. (H) Glucose-Citrate Communities with high-Overlap (O>0.98) have significantly higher mean Dissimilarity than Glucose-Glucose communities or Citrate-Citrate communities in the same Overlap range (O >0.98) (Welch Two-sample t-test).

**Table 1.** Specific taxa are strongly associated with high Dissimilarity in replicate habitats. For each of the 10 most commonly observed on glucose ESVs we performed one-tailed t-test to determine whether pairs of highly overlapping communities (O>0.98) both containing that ESV had higher Dissimilarity than pairs where at least one community did not contain the ESV. In this table, we report the identity of the ESV, the Number of communities in which that ESV is found (N) the difference in mean Dissimilarity, the T-statistic and the p-value (Bonferroni corrected for multiple comparisons). See Table S1 and S2 for a similar analysis applied to Leucine and Citrate Communities.

| ESV | N | Difference in Mean | T-statistic | P.value |
| --- | --- | --- | --- | --- |
| Citrobacter | 25 | 0.323111938 | 9.4851911 | 2.889193e-10 |
| Yersinia | 19 | 0.285958761 | 7.0293011 | 4.243258e-06 |
| Pseudomonas | 57 | 0.182202272 | 6.9293153 | 3.337966e-09 |
| Enterobacteriaceae | 38 | 0.240597425 | 6.4377452 | 8.930823e-07 |
| Pseudomonas.2 | 41 | 0.138617434 | 3.9956516 | 9.031112e-04 |
| Raoultella | 45 | 0.116423520 | 3.6636783 | 2.486771e-03 |
| Pseudomonas.3 | 18 | 0.006072537 | 0.1841094 | 1.000000e+00 |
| Pseudomonas.1 | 24 | -0.030734501 | -1.0303755 | 1.000000e+00 |
| Pseudomonas.4 | 27 | -0.035440920 | -1.2724340 | 1.000000e+00 |
| Klebsiella | 65 | -0.271582271 | -7.0743722 | 1.000000e+00 |

### Communities assembled in identical environments exhibit a negative correlation between Dissimilarity and Overlap

We first addressed the question of whether communities assembled in identical environments do indeed give rise to a negatively sloped DOC curve at high Overlap. To test this prediction, we took all pairs of communities that had been assembled in the same nutrient-limited habitats, by pooling together every possible pair of glucose communities, as well as every pair of citrate (and of leucine) assembled communities. We then measured the Dissimilarity and Overlap for each pair. Applying the same type of statistical analysis used in the original study by Bashan et al (Robust LOWESS regression; see Methods for details [18]), we find that the DOC (but not the controls; **Fig. S1**) does indeed exhibit a negative slope at high values of Overlap (**Fig. 1B**), and this is also confirmed by a standard linear regression (*D*=*D*_0_+*mO*; *m* = −0.56, p<0.002 by Bootstrapping; Methods) applied to the points with higher than median *O* (**Fig. 1B**, inset; Methods). The same is true when we analyze each of the three nutrient environments separately (**Fig.1C**), and it also holds when we separate those communities assembled from either the same or different inoculum (**Fig. 1D-E**). In contrast, a statistically significant negative correlation between Dissimilarity and Overlap is not observed for community pairs that are assembled in different environments (e.g. one in citrate medium, one in glucose medium) (**Fig. S2**).

An expectation of DOA is that communities that contain the same taxa in identical habitats should have them at highly similar equilibrium abundances, as the underlying population dynamics would be strongly similar. By contrast, communities assembled in different nutrient habitats are not expected to have similar species abundances even when they happen to share a large proportion of common species (high Overlap), as we do not necessarily expect their dynamical equations to be similar. Consistent with the first hypothesis, we find that, in identical nutrient conditions, the majority of our high-Overlap (O>0.98) communities have low Dissimilarity (**Fig. 1F,** see S1 for comparison to null) (Mean= 0.188, Median=0.132, IQR=0.23). To test the second hypothesis, we considered “mixed” pairs of communities, where each community in the pair was assembled in a different environment. For instance, we find that glucose-citrate pairs (which exhibit no correlation between *D* and *O* (**Fig. S2-3**)) have a similar overlap distribution to glucose-glucose and citrate-citrate pairs (**Fig.1G**). Yet, the distribution of Dissimilarities for these high-Overlap pairs (O> 0.98) is shifted up compared to glucose-glucose pairs (t =−12.79, p-value < 0.001, Welch two-sample test) and citrate-citrate pairs (t =−11.965, p-value < 0.001, Welch two-sample test) (**Fig.1H**). This shift persists even if we only consider communities assembled from the same inoculum (**Fig.S4**) and is robust to the Overlap threshold chosen (**Fig.S5**)).

This last finding is consistent with the idea that population dynamics and equilibria are strongly convergent when the environments are identical, but not necessarily when the environments are different. This lends support to the null assumption that species interactions with the environment and with each other are different in different environments, but strongly convergent in identical environments. Our results support the prediction that identical environments will generate a negative statistical correlation between *D* and *O*, whereas different environments will not.

### Specific taxa can be strongly associated with high Dissimilarity in replicate habitats

As can be visually appreciated in **Fig. 1**, when environments are identical and Overlap is high the Dissimilarity in our experimental communities is generally small. However there are numerous deviations from this rule, and we find multiple community pairs with high Overlap that still show high levels of Dissimilarity. Considering only communities with an Overlap > 0.98, we find that 12% of glucose Pairs, 9% of citrate pairs, and 17% of leucine pairs have 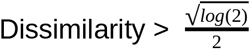 (which is half of the maximum possible Dissimilarity, calculated through the root Jensen–Shannon divergence) (**Fig. 1F**). Similar values are also obtained if we only consider communities assembled from the same inoculum, (14% of glucose pairs, 9% citrate pairs, 8% of leucine pairs) (**Fig. S4A**). In sum, we find that communities can be dominated by the same set of ESVs in identical habitats, yet these ESVs, may exist at very different abundances.

We set out to investigate whether communities that deviate from the average trend captured by the DOC could be associated with the presence of specific taxa. To that end, we first selected pairs of glucose communities with high Overlap (higher than the median of 0.98) that were assembled from the same initial pool of species (for a similar analysis in citrate or leucine, see Table S1 and S2). We then tested whether any of the 10 most commonly observed ESVs (corresponding to ESVs found in at least 18 of the 92 glucose communities), were statistically associated with high Dissimilarity. Six of these ESVs had higher Dissimilarity than expected by chance (Bonferroni Corrected p<0.05, one-tailed t-test). Of these, an ESV of the genus *Citrobacter* had the largest effect (Table S1).

To further investigate this point, we split all pairs of communities by whether both contain this *Citrobacter* ESV (group I), only one contains it (group II), or none does (group III) (**Fig. 2A**). The mean Dissimilarity is higher for group I than group II (0.427 vs 0.175, p<0.01, one-tailed t-test), and for group II than group III (0.175 vs 0.088,p<0.01, one-tailed t-test) **Fig. 2B**. As shown in **Fig. S6**, our results are robust to our choice of a “High-Overlap” threshold of 0.98. If *Citrobacter* ESV were indeed associated with alternative dynamical states (either through multistability, or through their contribution to alternative dynamical equations when they are part of the community), we would also expect the DOC to flatten for glucose-glucose communities that contain it, relative to those that do not. That is indeed the case, as shown in Figs. **2C-D**, see **Fig. S7** for controls).

**Fig. 2:**
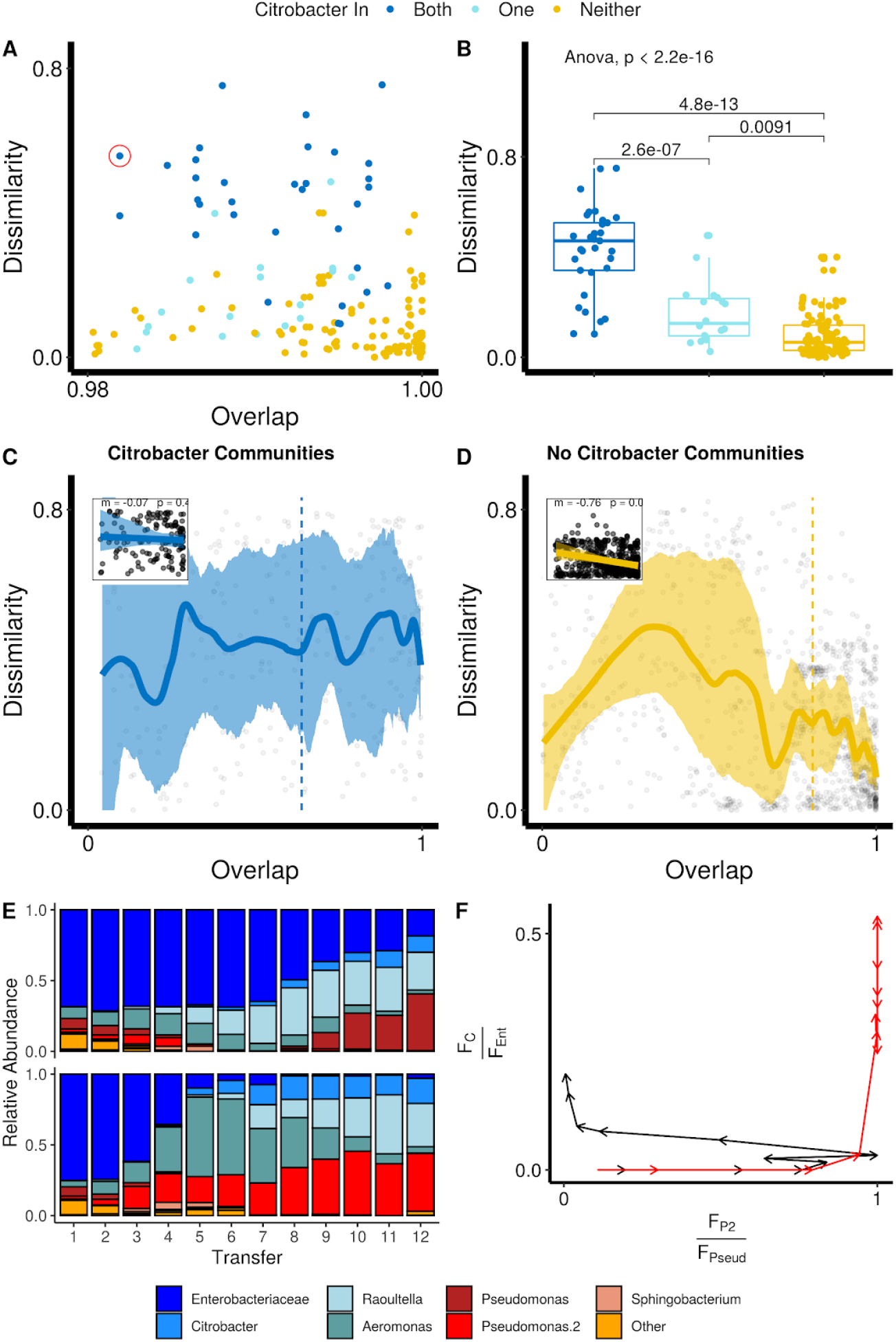
A *Citrobacter* ESV is associated with dynamical dissimilarity in communities assembled in replicate environments. (A) Dissimilarity and Overlap of microbial community pairs assembled from the same regional pool on M9 + Glucose with above-median Overlap (*O*>0.98). (B) Dissimilarity of the same set of communities. For (A) and (B) we label communities by whether *Citrobacter* ESV is found in both communities (dark blue), only in one community (light blue) or in neither community (yellow). (C) DOC of all pairs of microbial communities assembled on glucose that contain *Citrobacter* ESV (n = 25). (D) DOC of all pairs of microbial communities assembled on glucose that do not contain *Citrobacter* (n = 67). (E) Population dynamics for one pair of glucose communities with high Overlap (*O*=0.98) and high Dissimilarity (*D*=0.56) (highlighted in panel A with the red circle). Structure of the two communities at the ESVlevel at every transfer. (F) Phase portraits illustrate the dynamics of the most abundant Enterobacteriaceae and Pseudomonadaceae ESV within those two communities. *F_C_* and *F_P2_* represent the fraction of the *Citrobacter* and *Psuedomonas.2* ESVs in the population, whereas *F_Ent_* and *F_Pseud_* represent the fractions of the Enterobacteriaceae and Psuedomonoadacea Families. Dynamics are highly convergent until the third transfer, after which the communities diverge to alternative states.

Similar results were found for citrate communities and leucine communities, where we found a *Raoultella* ESV and a *Pseudomonas* ESV associated with higher Dissimilarity respectively (**Supp Table S1–S2**, and **Fig. S8**–**S9**). Our results thus reveal that the presence of a single ESV in a community may be strongly associated with alternative states in community assembly, even amongst communities that are assembled in the same environment from the same regional pool and contain highly overlapping sets of taxa.

## Discussion

The first part of this paper tests two fundamental predictions of DOA. First, using publicly available data from a recent experiment we show that community assembly in identical environments does lead to a negative correlation between Dissimilarity and Overlap and a negatively sloped DOC. Second, we show that, as expected, communities assembled in identical environments that also contain highly overlapping sets of taxa have them at strongly convergent abundances. This is consistent with what one would expect if their dynamical equations were identical. Likewise, we would not expect identical dynamical equations in different nutrient habitats. Consistent with this expectation, we found that communities assembled in different nutrient habitats do not have convergent species abundances, even when they happen to share a high number of taxa. This is reflected in the data by a shift in the Dissimilarity to higher values for those communities with very large Overlap.

Our results also indicate the existence of specific taxa that are associated with high Dissimilarity in replicate habitats. In glucose communities, we find an ESV of the genus *Citrobacter*, which not only predicts high Dissimilarity between community pairs when it is present in at least one of the communities, but it also flattens the DOC. Intriguingly, these results are in line with one of the main findings of the Bashan study, which was that microbiomes disrupted by *C. difficile* infection did not exhibit a negative correlation between Dissimilarity and Overlap, and Dissimilarity remained constant even as Overlap increased [18]. The negative DOC slope was recovered after a fecal microbiota transplantation, which restored a healthy microbiota and cured the disease. Although the reasons for this finding may be very different from the similar result found in our communities, our results indicate that a tight association between specific taxa and the flattening of the DOC may be seen even in the absence of an immune system or a complex host.

The publicly available data of Goldford et al [13] includes the measurement of the complete population dynamics for one glucose pair with high-Overlap and high Dissimilarity (highlighted in **Fig. 2A**). Both communities in the pair contain the *Citrobacter* ESV. For this pair, we find that the population dynamics are initially strongly convergent between the two communities, but bifurcate after ~20 generations and subsequently diverge into alternative compositions(**Fig. 2E-F**), suggesting the potential presence of true multi-stability (i.e., multiple stable steady states are associated with the same set of species). Of course, other possibilities exist. For example, the population dynamics are just not identical even in replicate habitats due to the accidental violation of the second assumption of DOA. Moreover, the seemingly high overlap of the two communities at the ESV level might not reflect their differences at the stain level due to rapid evolution. *Citrobacter* may also have more sensitive interactions with the rarer members of the community. It is important to consider that environments do not just passively select for species and determine their interactions, but rather they are dynamically shaped by the taxa growing in them [21–24]. Therefore, it is possible that although the supplied nutrients are the same in two communities, the environment experienced by the members of our communities is actually different, through the different effects that species have on it [21].

One limitation of our study is that the communities investigated are species-poor, and many of our community pairs will only share a few species in common. This can be potentially problematic because it may affect the Dissimilarity measurement [18]. Despite this caveat, the overall good agreement between our findings and the expectations of DOA suggests that species richness is not necessarily an impediment for the application of DOA to taxonomically poor natural communities. Further research would be needed to establish the precise conditions under which this would be true. More generally, others have argued that communities assembled along an environmental gradient may also give rise to negative correlations between Dissimilarity and Overlap [19]. It would be important to test this prediction experimentally, but this falls beyond the scope of this paper.

It is thus important to remark that although sets of communities assembled in identical habitats present a negatively sloped DOC, the reverse statement is not necessarily true: the presence of a negatively sloped DOC does not necessarily mean that the habitats are identical to each other. In fact, grouping together all communities in the experiment, including those assembled in identical and different environments and projecting them all into the same Dissimilarity-Overlap plot, we find a negative correlation between *D* and *O* and a negatively sloped DOC (**Fig. S10**). The reason is that the strong effect of identical habitats overpowers the effect of non-identical ones. In our case, we know the environmental factor that was critically different among these habitats (the single limiting nutrient), but this is not something that is trivial to identify in any given natural habitat, even if we remove any known factors of variation across habitats.

Notwithstanding these important caveats, our results confirm in a controlled set of experiments that microbial dynamics in replicate habitats are strongly convergent on average and lead to a negatively sloped DOC. In recent years, negatively sloped DOC has been detected for the microbial communities assembled in some (but not all) human body sites [18], as well as for mycorrhizal fungal communities [20]. In order to correctly interpret these results, it is critical to benchmark the technique not only against simulations, but also against experimental communities whose assembly is well understood. Our results have provided a first empirical benchmark using well-controlled communities. We hope that these findings will contribute to grounding the expectations and intuitions behind DOA, and contribute to its application to microbial communities assembled in natural environments.

## Methods

### Community Assembly Experiment

We analyzed publicly available data from a recent set of *in-vitro* microcosms experiments [13]. Briefly, diverse microbial communities were isolated from natural ecosystems and used as the inoculum for a batch culture containing M9 + one of three carbon sources (Glucose, Citrate, Leucine). Cultures were passaged every 48 hrs with a dilution factor of 125x and after each transfer, a sample was taken and stored for 16S community sequencing (**Fig. 1A**). The experiment was conducted for 12 transfers by which point communities appear to have reached a stable population equilibrium. In total, 276 communities were allowed to self assemble (7-8 Replicates per Inoculum and 12 Inoculum per carbon source). Community structure was determined at the end of the twelfth growth period for all communities using 16s rRNA amplicon sequencing. A subset of communities was sequenced at each transfer allowing community structure to be tracked through time.

### Calculating Dissimilarity and Overlap for community pairs

To account for differences in sequencing depth we first normalize all communities so that each community is represented by the same number of sequences. Briefly, for each community we create a sample community of *N* reads sampled from the original read pool without replacement. Here *N=4397* was used as this is the minimum number of reads for all communities analyzed. We calculated the relative abundance of each Exact Sequence Variant (ESV) in all communities. For each pair of communities, we follow Bashan et al [18] and calculate Overlap and Dissimilarity between relative abundance vectors *x* and *y*. Overlap is given by

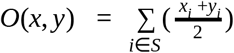

Where S is the set of ESVs found in both communities. For the calculation of Dissimilarity only the shared ESVs are considered, and the relative abundance of shared taxa is renormalized to add up to 1. The Dissimilarity between renormalized vectors *x* and *y* can be calculated as root Jensen-Shannon divergence

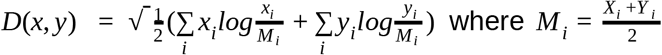

We calculated the Overlap and Dissimilarity for every pair of our 276 communities at transfer 12 (a total of 37950 pairwise comparisons). From this dataset, any pairwise comparison in which the communities shared fewer than 2 taxa in common is excluded. These were removed as for these pairs D(*x*,*y*) is always 0. In total this gave us a dataset of 23592 pairwise comparisons.

### Fitting DOCs

In Bashan et al [18] estimated using the Robust LOWESS a non-parametric scatterplot smoothing method. When analyzing all pairs of communities (**Fig. 1B**, **Fig.S2A**, **Fig2C-D**), we implement the same method using the LOESS function from R default stats package with the following parameters (span=0.2, family=“symmetric”, iterations=5). To compare slopes across different subsets of pairs (Insets **Fig. 1B**, **Fig. 1C-E**, **Fig.S2B-D**, Insets **Fig.2C-D)** we use a simple OLS regression on data points with above-median Overlap as was also done by Bashan at et al [18] and later by Kalyuzhny and Shnerb [19].

### Estimating Confidence Intervals and P-value for DOC

We implemented the same bootstrapping algorithm used by Bashan et al [18]. We repeat this bootstrap algorithm 500 times and repeated all our analysis on every bootstrap realization. Confidence Intervals in **Fig.2** and **Fig.4** represent 95% percentiles of the curves fitted to the bootstrapped data. The reported p-values for the regression slopes (*m*) represent the fraction of bootstrap realization for which the OLS slope is negative (main text and Fig.2, Fig.4, Fig.S3).

### Randomized Data

Each time a DOC is shown, we repeat the analysis on a randomized dataset in which species assemblage and abundance distribution are kept but the abundance of each taxon is randomized following Bashan et al [18]. The randomized results are shown in **Fig. S1**, **Fig.S3**, and **Fig. S7**.

### Code Availability

All the analysis was conducted R. The R Code and datasets analyzed are available at https://github.com/vilacelestin/VilaLiuSanchez2020

## Supplementary Figures

**Supplementary Figure 1:**
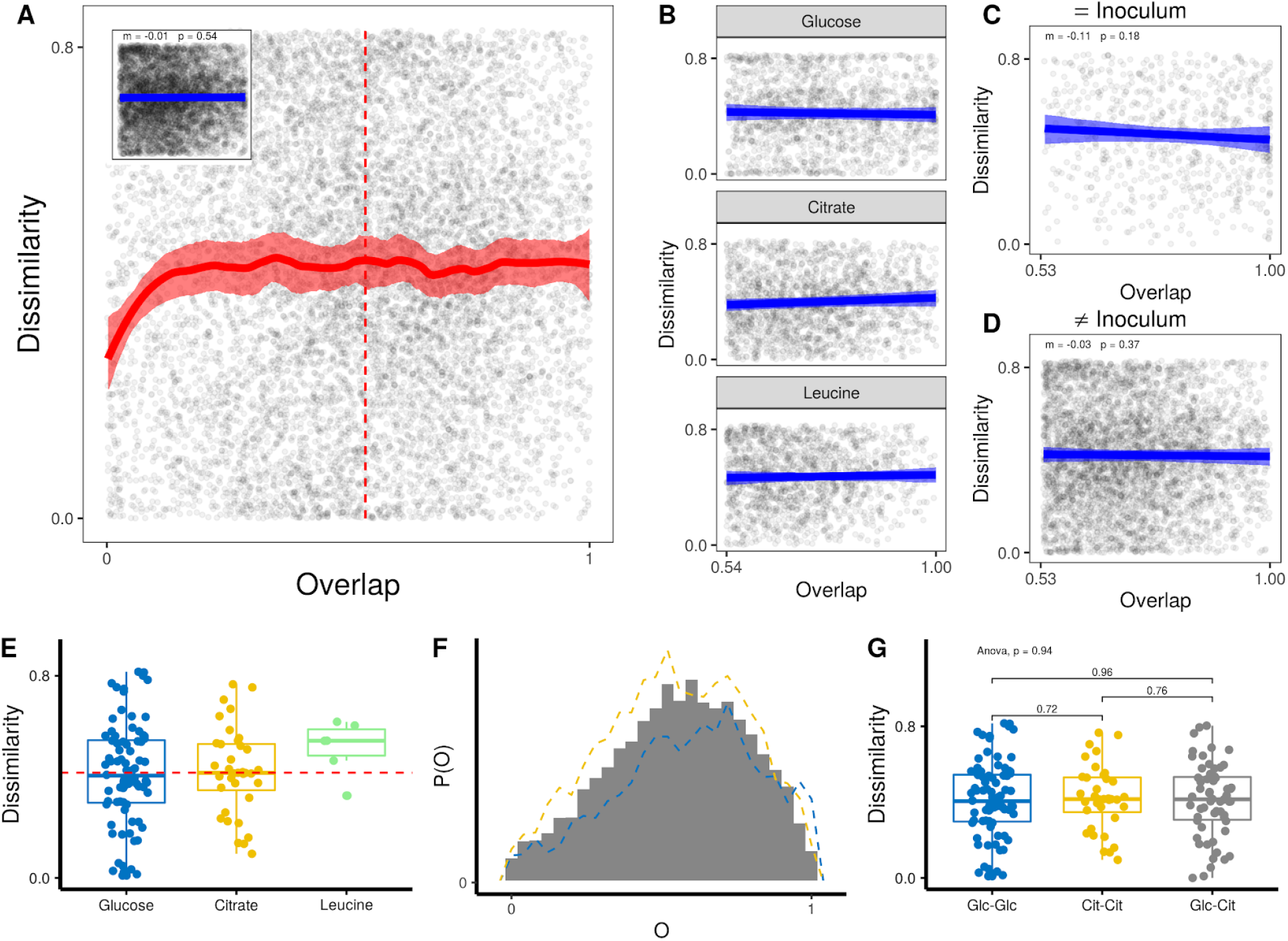
A statistically significant negative correlation between Dissimilarity and Overlap is not observed for randomized samples (see Methods). Panel A is a randomized version of **Fig. 1B**. Panel B is a randomized version of **Fig. 1C**. Panel D is a randomized version of panel **Fig. 1E.** Panel E is a randomized version of **Fig. 1F**, Panel F is a randomized version of **Fig. 1G**. Panel G is a randomized version of **Fig. 1H**.

**Supplementary Figure 2.**
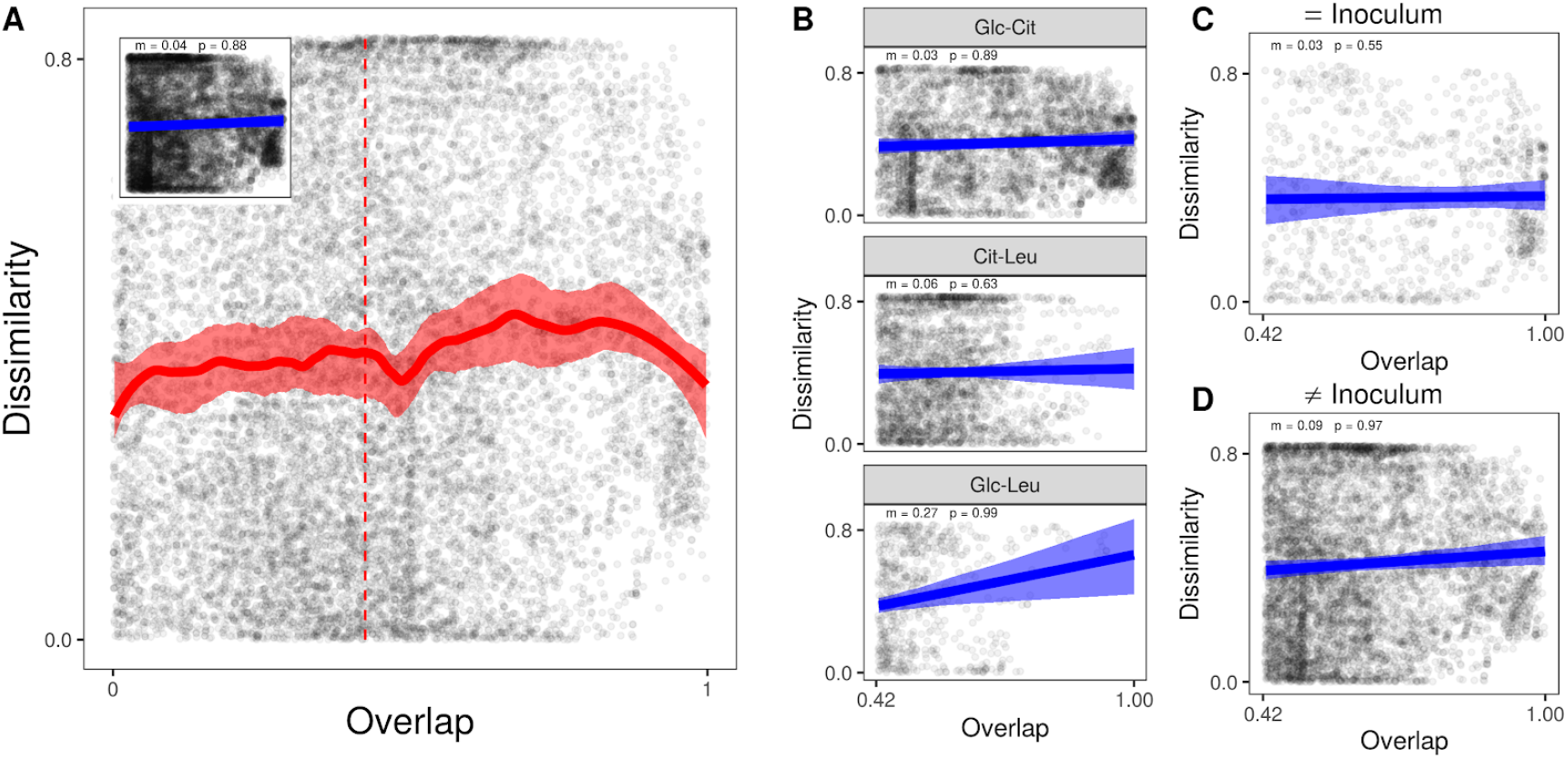
A statistically significant negative correlation between Dissimilarity and Overlap is not observed for community pairs that are assembled in different environments (e.g. one in citrate medium, one in glucose medium). This holds true regardless of whether we consider: (A) all pairs in different environments; (B) Each contrasting environment separately. (C) Community pairs assembled from the same inoculum; or (D) Community pairs assembled from different inocula.

**Supplementary Figure 3:**
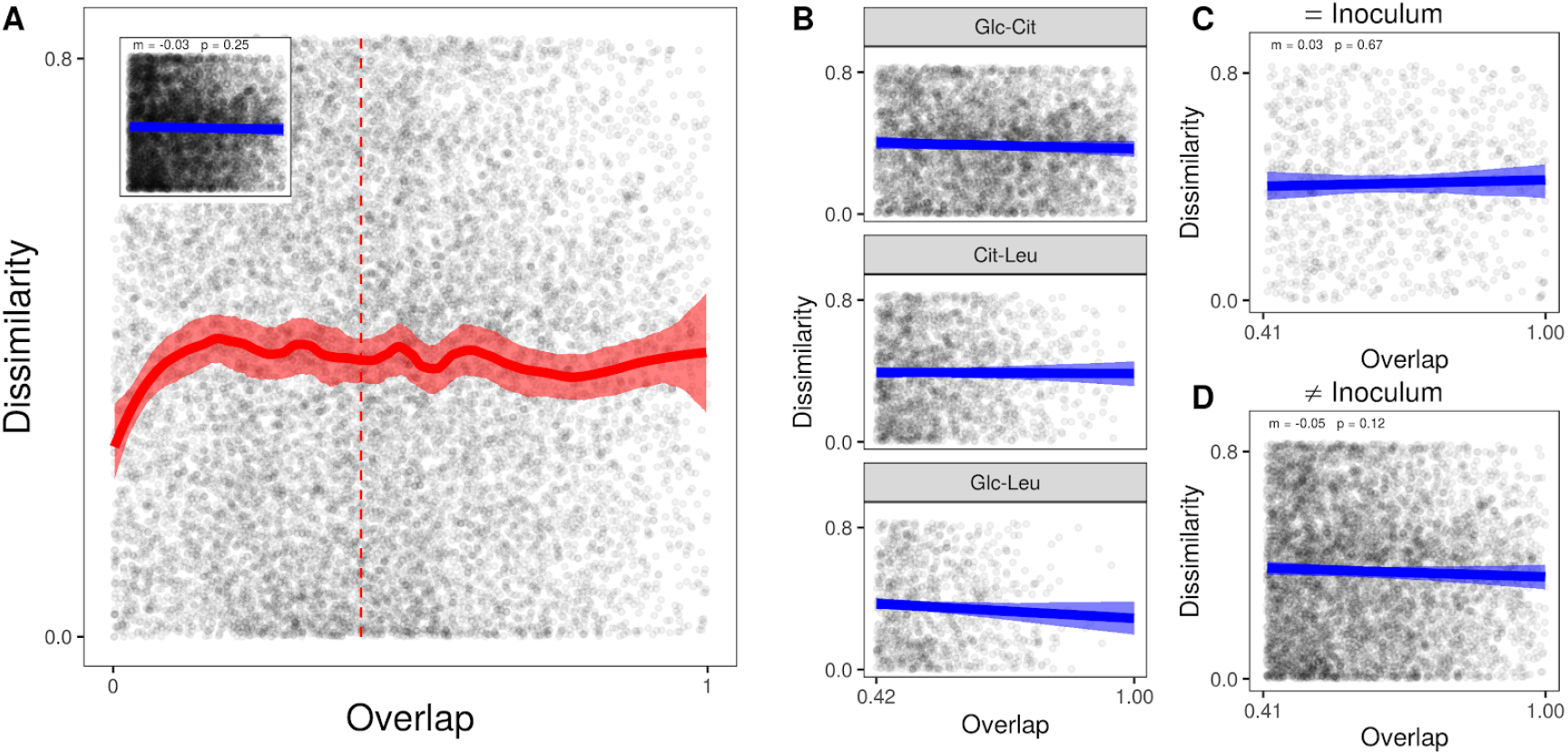
Randomized Controls for Figure S2 also do not show a negative correlation. Panel A,B C and D show the randomized controls for Figure S2 A,B,C and D respectively.

**Supplementary Figure 4:**
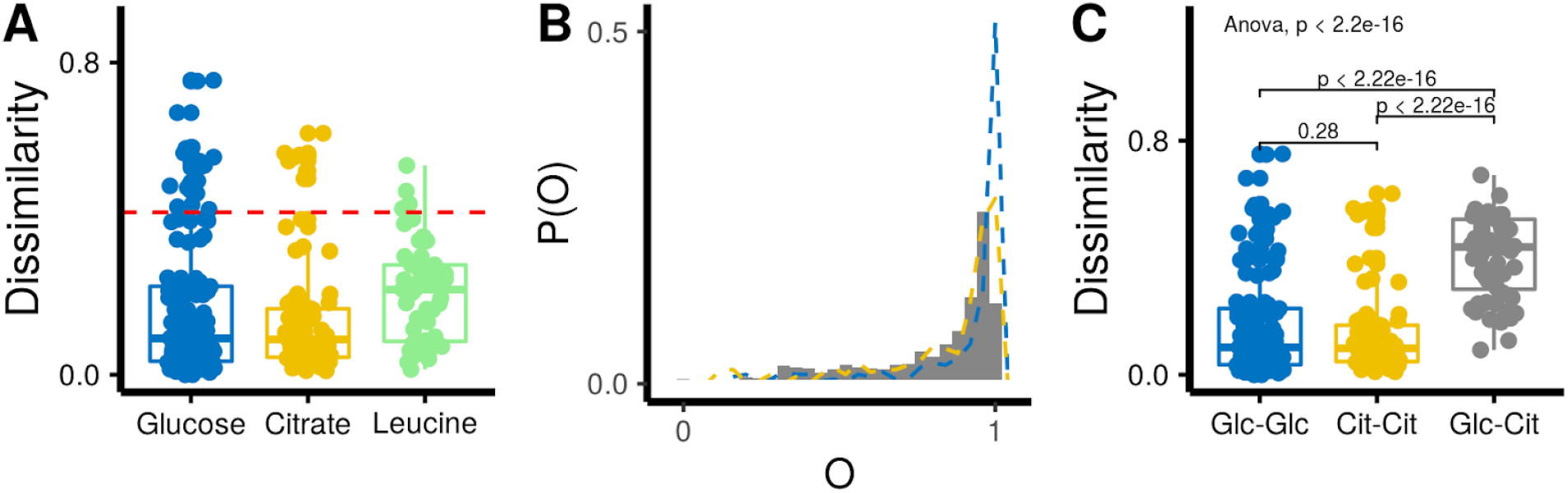
Reproduction of Figure 1 F-H considering communities that come from the same inoculum. (*A*) Distributions for community pairs assembled in the same environment from the same inoculum (both in glucose, both in citrate, or both in leucine) with high overlap (O>0.98). The dotted red-line is at half the maximum possible 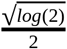. (B) Histogram showing distributions of Overlaps for community pairs assembled from the same inoculum where one has been assembled on glucose and the other on citrate. The dotted lines give the frequency polygon for Glucose-Glucose Community pairs and Citrate-Citrate community pairs (Blue and yellow respectively). We use the same binwidth (.04) for both histograms and frequency polygons so the two are comparable. (C) Glucose-Citrate Communitie from the same Inoculum with high-Overlap (O>0.98) have significantly higher mean Dissimilarity than Glucose-Glucose communities or Citrate-Citrate communities in the same Overlap range (O >0.98) (Welch Two-sample t-test).

**Supplementary Figure 5:**
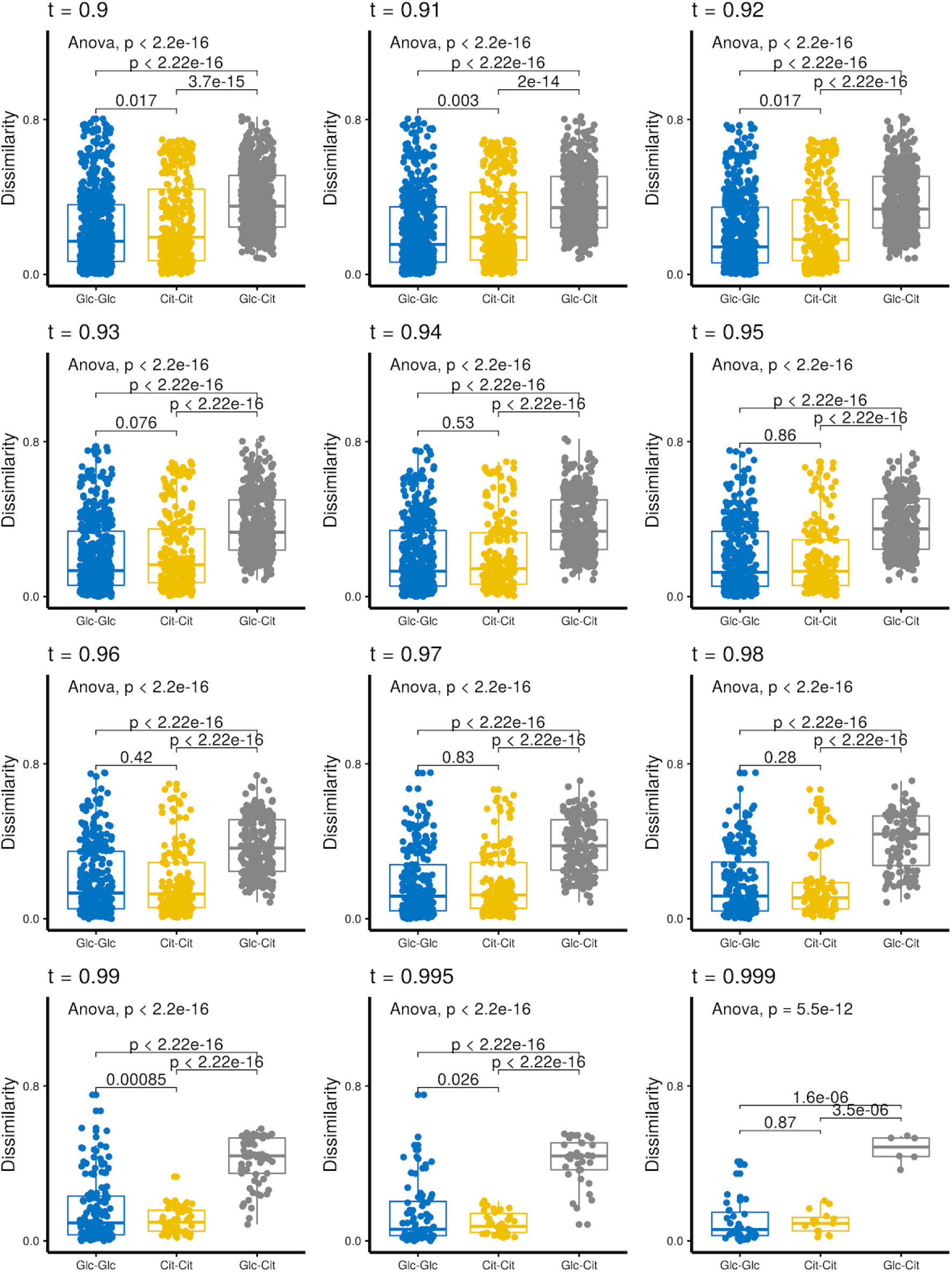
The results plotted in Figure 1H are robust to the exact choice of threshold Overlap (*t*). In the main text, we show the result for *t* = 0.98, whereas here we give the results for a wide range of thresholds (*t*)

**Supplementary Figure 6:**
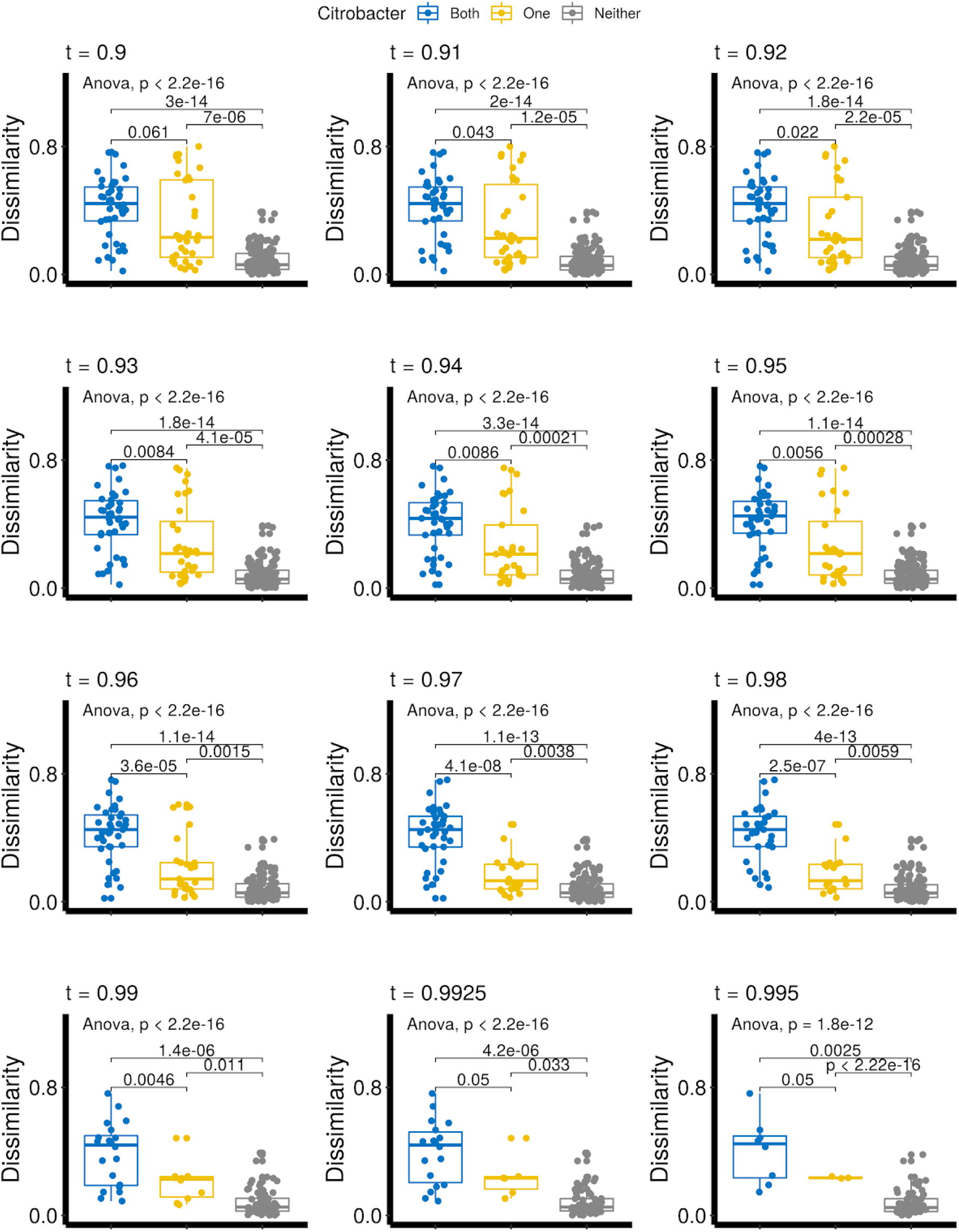
The results plotted in Figure 2B are robust to the exact choice of threshold Overlap (*t*). In the figure, we show the result for the median overlap, whereas here we give the results for a wide range of thresholds (*t*).

**Supplementary Figure 7:**
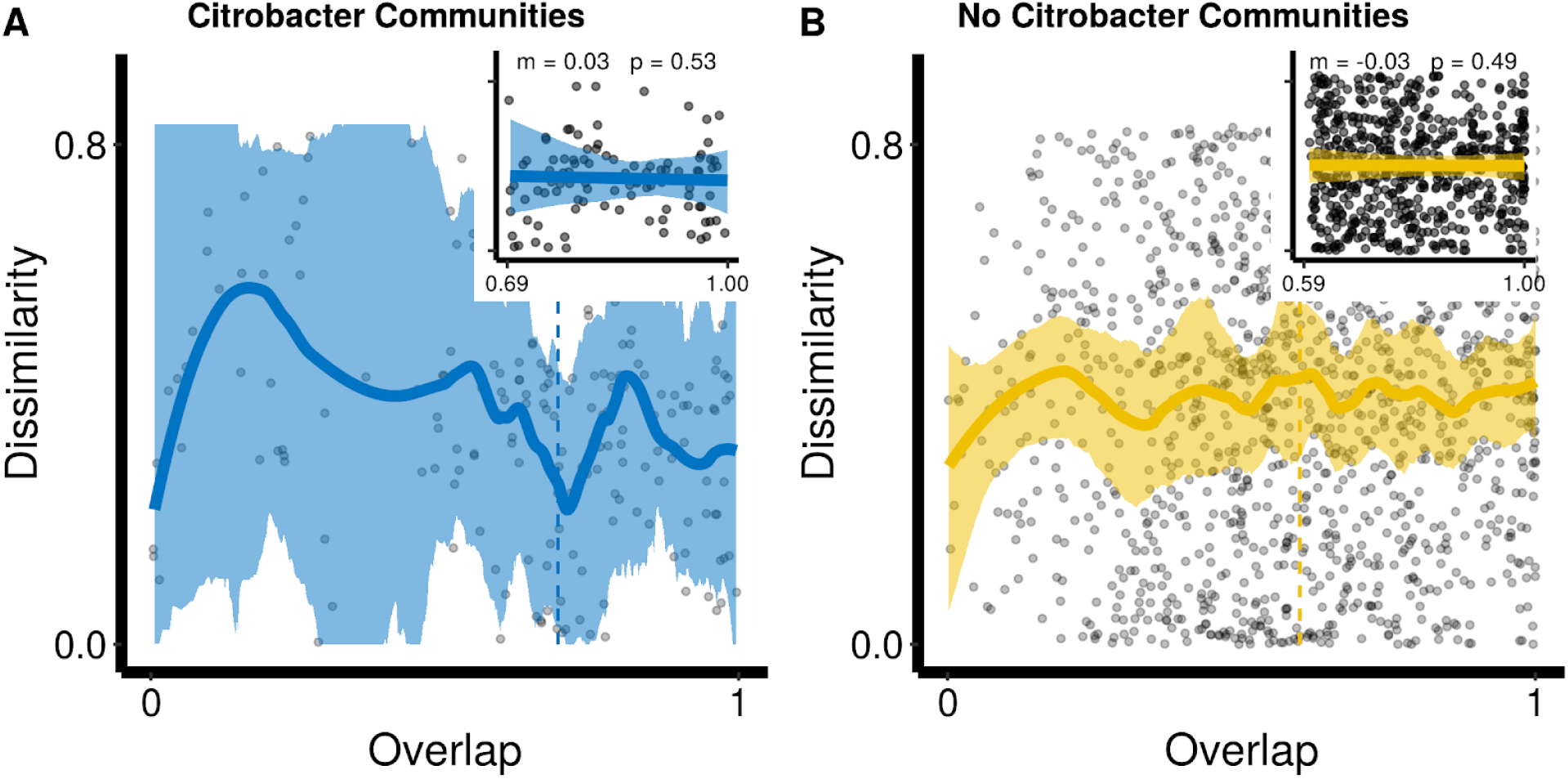
Communities assembled in glucose which contain (A) or do not contain (B) *Citrobacter* ESV do not display a negative DOC nor a statistically significant negative regression coefficient (insets) between Dissimilarity and Overlap when the ESV abundances are randomized (Methods).

**Supplementary Figure 8:**
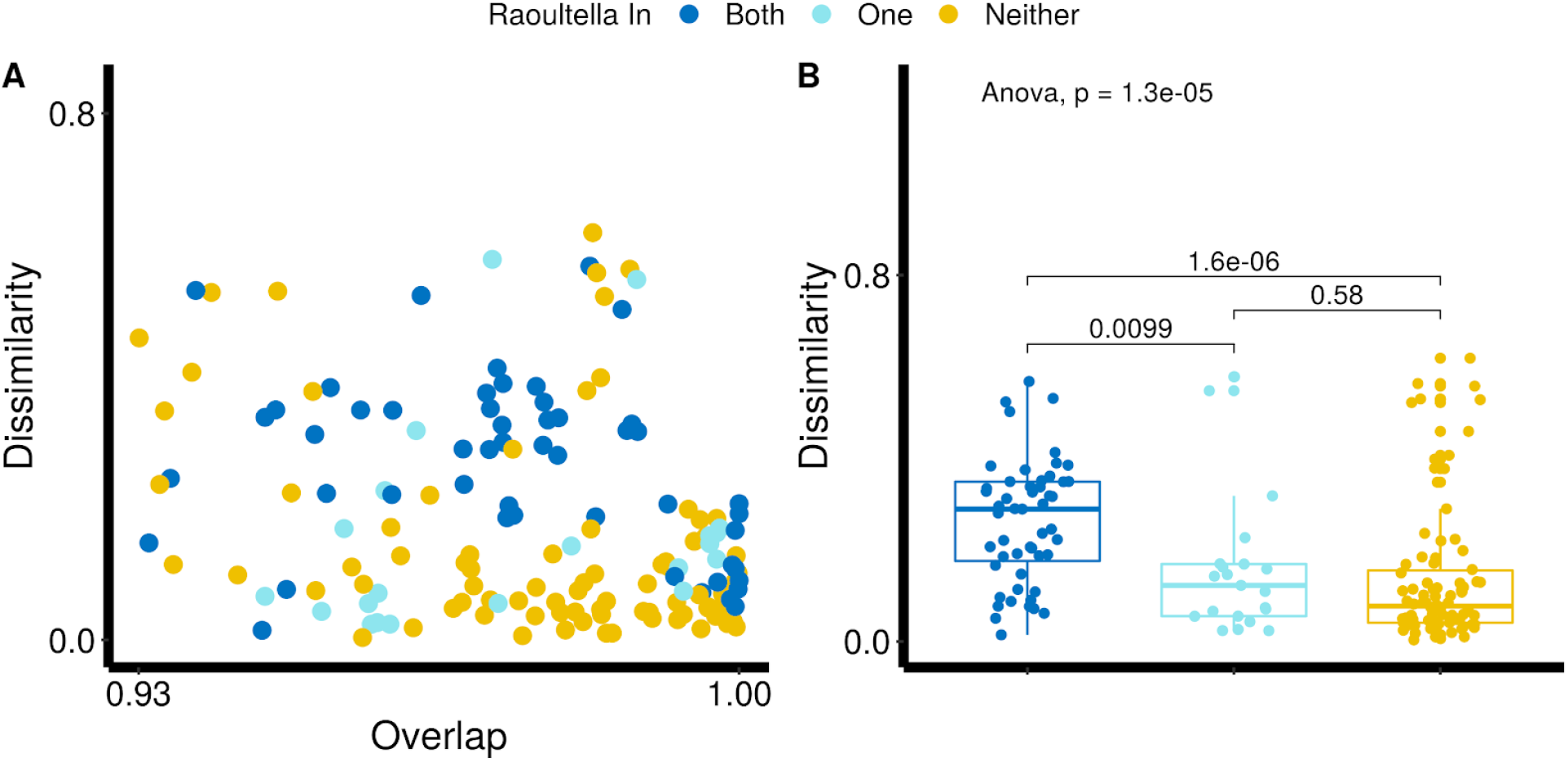
A *Raoultella* ESV is associated with dynamical dissimilarity in community assembled on Citrate. Dissimilarity and Overlap of microbial community pairs assembled from the same regional pool on M9 + Citrate with above-median Overlap (*O*>0.93) (B) Dissimilarity of the same set of communities. For (A) and (B) we label communities by whether *Raoultella* ESV is found in both communities (dark blue), only in one community (light blue) or in neither community (yellow).

**Supplementary Figure 9:**
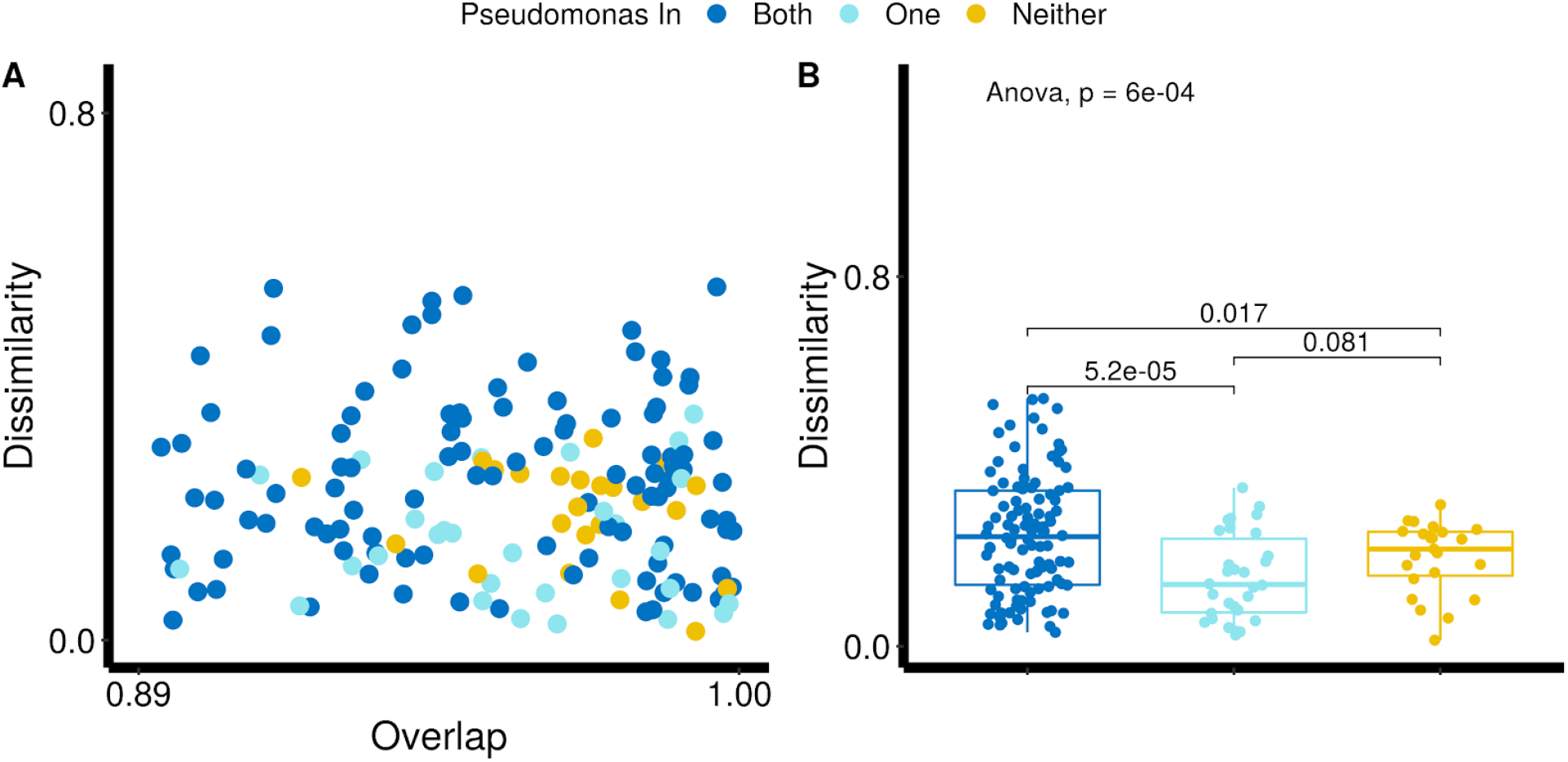
A *Pseudomonas* ESV is associated with dynamical dissimilarity in community assembled in replicate environments. Dissimilarity and Overlap of microbial community pairs assembled from the same regional pool on M9 + Leucine with above-median Overlap (*O*>0.89) (B) Dissimilarity of the same set of communities. For (A) and (B) we label communities by whether *Citrobacter* ESV is found in both communities (dark blue), only in one community (light blue) or in neither community (yellow).

**Supplementary Figure 10:**
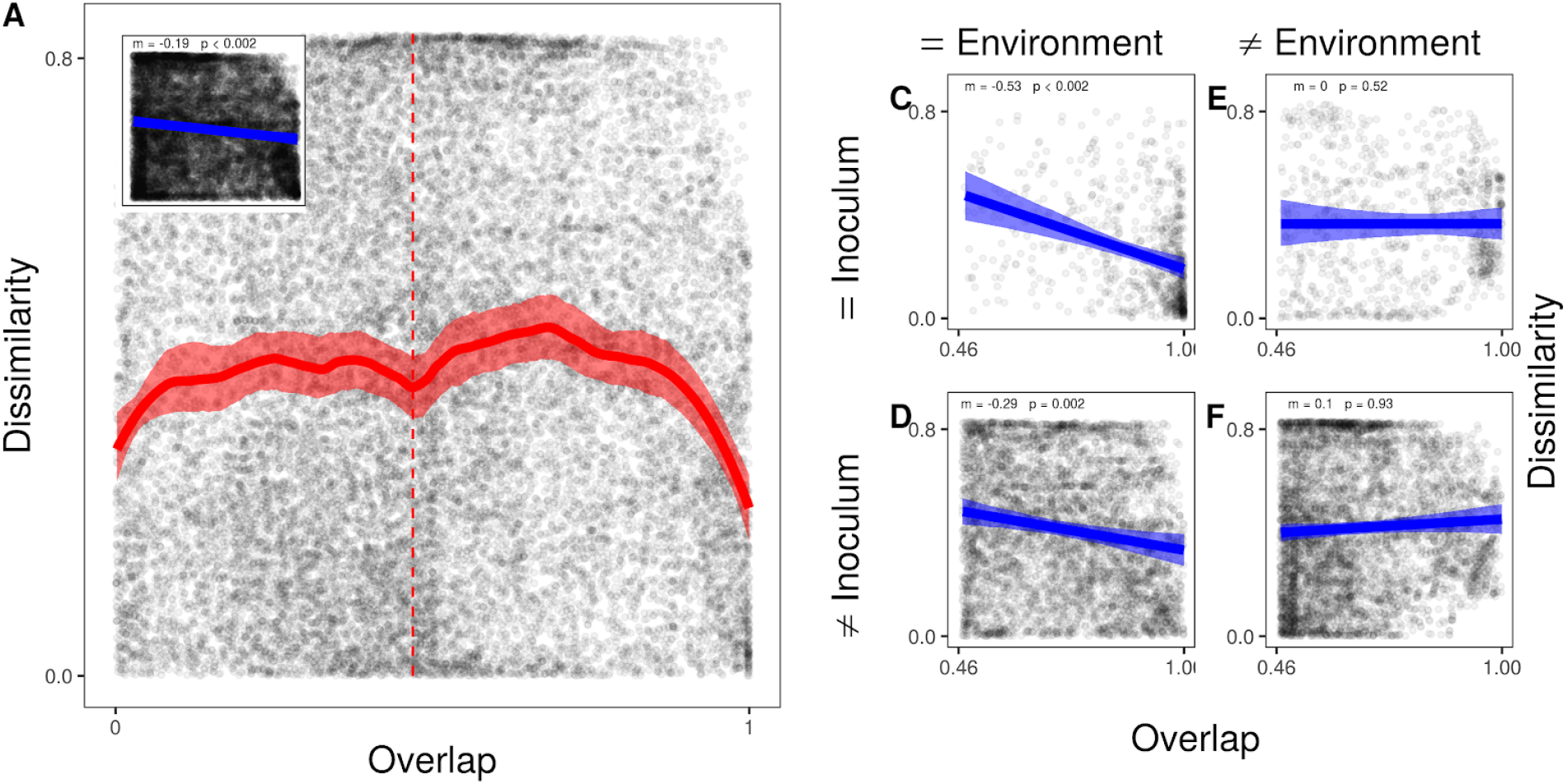
DOC of all microbial community pairs analyzed in this study. (n = 276 Samples). Shaded regions indicate the 95% confidence interval (Methods). We do observe a negative DOC but this is largely driven by community pairs assembled in the same environment. (Panel C and D).

**Supplementary Table 1:** Repeat of analysis in Table 1 for Citrate communities.

| ESV | N | Difference in Mean | T-statistic | P.value |
| --- | --- | --- | --- | --- |
| Raoultella | 35 | 0.12629694 | 5.2286408 | 4.422734e-06 |
| Ochrobactrum | 35 | 0.12322066 | 3.3623846 | 9.485359e-03 |
| Stenotrophomonas | 25 | 0.11614973 | 3.0643742 | 3.146488e-02 |
| Pseudomonas.1 | 52 | 0.07306871 | 3.0078636 | 1.540008e-02 |
| Bordetella | 33 | 0.07194774 | 1.5731749 | 6.666206e-01 |
| Enterobacteriaceae | 36 | 0.01868913 | 0.4511268 | 1.000000e+00 |
| Pseudomonas | 76 | 0.01021339 | 0.3730456 | 1.000000e+00 |
| Pseudomonas.2 | 34 | 0.00980748 | 0.2775113 | 1.000000e+00 |
| Klebsiella | 64 | -0.04085263 | -1.1243570 | 1.000000e+00 |
| Enterococcus | 28 | -0.04673677 | -1.7413145 | 1.000000e+00 |

**Supplementary Table 2.** Repeat of analysis in Table 1 for Leucine communities.

| ESV | N | Difference in Mean | T-statistic | P.value |
| --- | --- | --- | --- | --- |
| Pseudomonas | 74 | 0.0787870444 | 4.38572493 | 0.0001135545 |
| Delftia | 72 | 0.0511280633 | 2.55949503 | 0.0588186659 |
| Pseudomonas.8 | 25 | 0.0697182320 | 2.45727588 | 0.1074751384 |
| Pseudomonas.9 | 29 | 0.0545401992 | 1.64290966 | 0.5656843823 |
| Aeromonas | 25 | 0.0187680529 | 0.76539029 | 1.0000000000 |
| Stenotrophomonas | 70 | 0.0042829219 | 0.21288729 | 1.0000000000 |
| Comamonas | 30 | -0.0003044766 | -0.01618936 | 1.0000000000 |
| Bordetella | 33 | -0.0061502179 | -0.31991060 | 1.0000000000 |
| Pseudomonas.1 | 55 | -0.0180839614 | -0.88178585 | 1.0000000000 |
| Pseudomonas.3 | 28 | -0.0272273747 | -1.33162337 | 1.0000000000 |

